# Nicotinamide riboside kinase-2 inhibits JNK pathway and limits dilated cardiomyopathy in mice with chronic pressure overload

**DOI:** 10.1101/2021.10.14.464350

**Authors:** Syeda Kiran Shahzadi, Hezlin Marzook, Rizwan Qaisar, Firdos Ahmad

## Abstract

Nicotinamide riboside kinase-2 (NRK-2) has recently emerged as a critical regulator of cardiac remodeling however, underlying molecular mechanisms is largely unknown. To explore the same, *NRK2* knockout (KO) and littermate control mice were subjected to trans-aortic constriction (TAC) or sham surgeries and cardiac function was assessed by serial M-mode echocardiography. A mild cardiac contractile dysfunction was observed in the KOs at the early adaptive phase of remodeling followed by a significant deterioration during the maladaptive cardiac remodeling phase. Consistently, *NRK2* KO hearts displayed increased cardiac hypertrophy and heart failure reflected by morphometric parameters as well as increased fetal genes *ANP* and *BNP* expressions. Histological assessment revealed an extensive left ventricular (LV) chamber dilatation accompanied by elevated cardiomyopathy and fibrosis in the KO hearts post-TAC. In a gain-of-function model, NRK-2 overexpressing in AC16 cardiomyocytes displayed significantly attenuated fetal genes *ANP* and *BNP* expression. Consistently, NRK-2 overexpression attenuated angiotensin II-induced cardiomyocyte death. Mechanistically, we identified NRK-2 as a regulator of JNK MAP kinase and mitochondrial function where NRK-2 overexpression in human cardiomyocytes markedly suppressed the angiotensin II-induced JNK activation and mitochondrial depolarization. Thus, our results demonstrate that NRK-2 plays protective roles in pressure overload-induced dilatative cardiac remodeling and, genetic ablation exacerbates dilated cardiomyopathy, interstitial collagen deposition, and cardiac dysfunction post-TAC due, in part, to increased JNK activation and mitochondrial dysfunction.

## Introduction

A significant therapeutic advancement has been made to combat various chronic diseases such as cancer, however, heart failure (HF) still awaits an effective treatment strategy. The treatment for HF remains elusive, partly because the molecular mechanisms driving it are poorly understood. Cardiomyopathies (CM) including hypertrophic (HCM) and dilated cardiomyopathy (DCM) are among the leading causes of HF. DCM is characterized by enlargement of the left ventricular (LV) chamber and is often associated with dilatation of all four cardiac chambers in the chronic stage of disease [1]. Although cardiac contractile functions are minimally affected in the early stage of DCM, a reduced systolic function is observed in the advanced DCM. Unlike DCM, HCM is associated with LV wall thickening followed by a narrowing of the LV chamber, resulting in LV diastolic dysfunction [2]. Amongst all types of cardiomyopathies, DCM is the most prevalent cause of HF. The prevalence of DCM is infrequent in children and increases with advancing age, with higher frequency among men [3, 4]. No definite molecular targets are known to treat DCM and cardiac transplantation remains the only viable therapeutic option [5].

NRK-2 was first identified as a β1 integrin-binding protein, which regulates skeletal myoblast differentiation [6]. The pioneering study showed that overexpression of NRK-2 attenuated the fusion and terminal differentiation of the myoblast. Recently, we and others have identified that NRK-2 is required to protect pathological consequences under cardiac stressors [7-9]. Increased activation of P38 mitogen-activated protein kinase (MAPK) was, in part, responsible for the detrimental phenotype of NRK-2 deficient mice post-cardiac ischemia [8]. However, the role of NRK-2 in chronic pressure-overload (PO)-induced cardiac remodeling remains unknown.

NAD+ homeostasis disruption due to mitochondrial dysfunction is reported in numerous HF models including myocardial infarction (MI), PO and angiotensin II (Ang II) infusion models [10-14]. NRK-2 phosphorylates nicotinamide riboside (NR) to nicotinamide mononucleotide (NMR), which is then converted to NAD+ through nicotinamide mononucleotide adenylyl transferases (NMNAT). In several cardiac injury models, NRK-2 expression is found to be upregulated robustly [7], while NAMPT enzyme is found to be repressed. Hence, it was suggested that in a failing heart where the NAMPT pathway is inhibited, the NRK-2 pathway presents a common adaptive mechanism for NAD+ synthesis.

A variety of cardiomyocyte signaling pathways are regulated upon hemodynamic PO including G-protein coupled receptors (GPCRs), janus kinase/signal transducer and activator of transcription (JAK/STAT), phosphoinositide 3-kinase (PI3K) and MAPKs pathways such as extracellular regulated kinase (ERK), P38 and jun N-terminal kinase (JNK) [15-17]. Mostly both biomechanical and humoral factors are responsible for the activation of these pathways however, some are explicitly regulated by humoral factors. Persistent hemodynamic PO exerts cardiac mechanical stress which induces the release of factors such as Ang II, endothelin-1, and transforming growth factor-β, which further induces pathological cardiac hypertrophy and interstitial fibrosis [17, 18].

MAPK pathway is among the most prevailing signaling pathways engaged with cardiac hypertrophy and HF. Activation of each class of MAPKs seems to require a particular stimulus. For instance, activation of ERK1/2 is strongly observed in the presence of phorbol myristate acetate (PMA) and growth factors; however, their impact on JNK and p38 kinase is weak [19]. Similarly, in our previous study, we observed activation of p38 post-MI in NRK-2 deficient mice, but the activation of other MAPKs was unaffected [8]. Therefore, it is clinically important to investigate the role of NRK-2 in chronic PO-induced adverse cardiac remodeling and cardiomyocyte signaling.

Herein we show, for the first time, that NRK-2 is a critical regulator of JNK MAPK and dilated cardiomyopathy in a PO model. Employing *NRK2* null mice, we report that NRK-2 deficiency exacerbates LV chamber dilatation and cardiac dysfunction post-TAC due to increased cardiomyopathy and excess collagen deposition. These adverse cardiac phenotypes potentially caused due to excess JNK activation in the *NRK2* KO heart post-TAC. We strongly believe that these studies unravel the specific roles of NRK-2 in the chronic PO-induced maladaptive dilated cardiac remodeling.

## Method

### Generation of *NRK2* knockout mice

*NRK2* knockout (KO) mouse was purchased from The Jackson Laboratory (stock# 018638) which now discontinued from The Jackson Laboratory and embryonic stem (ES) cells are currently maintained by Knockout Mouse Phenotyping (KOMP) repository at UC-Davis (Stock # 059444-UCD). The detailed method of the generation of *NRK2* KO mouse was previously described [8]. Briefly, *NRK2* gene expression was disrupted by inserting ZEN-UB1 Velocigene cassette. The construct positive embryonic stem (ES) cells were injected into B6(Cg)-*Tyr*^*c-2J*^/J blastocysts. The C57BL/6NJ females mice were crossed with chimeric male mice followed by crosses with B6N.Cg-Tg(Sox2-cre)1Amc/J to remove neo cassette and with C57BL/6NJ to remove the *cre*-expressing transgene. Mice were maintained in heterozygous conditions (*NRK2+/-*) on the C57BL/6 background. The Institutional Animal Care and Use Committee of Vanderbilt University Medical Center approved all the animal procedures and treatments. All the animal-related experiments were performed at Vanderbilt University Medical Center.

### Trans-aortic constriction surgery in mice

The PO model was generated by trans-aortic constriction (TAC) as described previously [20, 21]. Briefly, 8-10 weeks old male *NRK2* KO and littermate control mice (n=16-17) were anesthetized through intraperitoneal injection of ketamine (50 mg/kg) and xylazine (2.5 mg/kg). The sleeping mice were intubated and the trachea was exposed by making an incision over the cervical midline. Mice were then intubated using a 20G needle which was connected to a ventilator, and the tidal volumes and respiration rates were adjusted according to the animal body weight as described previously [22, 23]. The aortic arch area was cleaned and constriction was performed by placing a 27-gauge needle over the aortic arch and tying it with a 7-0 nylon suture. The needle was carefully removed to generate a constriction of approximately 0.4 mm in diameter. In sham group animals, all the procedures were performed except knotting the aortic arch.

### Echocardiography

Cardiac contractile function and dimensions were measured in mice (n=9-14) through echocardiography as described previously [24, 25]. Briefly, transthoracic two-dimensional motion mode-echocardiography was performed at 0, 2, 4 and 6 weeks post-TAC using a 12-MHz probe (VisualSonic, Vevo2100). During echocardiography mice anesthesia was maintained by supplying oxygen inhalation containing 1.5% isoflurane. LV end-diastolic interior dimension (LVID;d) and end-systolic interior dimension (LVID;s) was measured and ejection fraction (EF) and fractional shortening (FS) values were calculated using the Vevo2100 program.

### Histochemistry and measurement of cardiomyocyte area

Histochemistry and cardiomyocyte size measurement were performed as described previously [26, 27]. Briefly, 6 weeks post-TAC (n=6 each) or sham (n=3 each) surgeries, mice were anesthetized and the whole heart was harvested and fixed using 4% paraformaldehyde. The cardiac tissues were dehydrated through serial concentrations of ethanol followed by embedding in paraffin. The longitudinal heart sections (5μm thick) were sliced and stained using Masson’s trichrome kit (Sigma-Aldrich# HT15-1KT) following the instruction manual. A 0.8X objective of a Nikon Eclipse 80i microscope was used to capture the whole heart image and, a 20X objective was applied to take five images from the LV region of the heart. Cardiomyocyte cross-sectional area was measured and analyzed in a blinded manner using NIS Element software.

### Determination of *in vivo* myocardial apoptosis

Cardiomyocyte apoptosis levels in 6 weeks post-TAC hearts was determined using in situ Cell Death Detection kit, TMR red (Roche# 12156792910) through terminal deoxynucleotidyl transferase-mediated dUTP nick-end labeling (TUNEL). The heart sections (n=3 each sham group, n=6 each TAC group) were treated as per kit manual recommendation and co-stained with α-actinin (Sigma# A7811, 1:200 dilution), a cardiomyocyte-specific marker, and 4,6-diamidino-2-phenylindole (DAPI). Sections were treated with mounting media and covered with a glass cover. A 20X objective of the Nikon Eclipse 80i fluorescence microscope was used to capture five random images of TUNEL-positive cardiomyocytes from the LV region of each longitudinal heart section. Quantification of TUNEL-positive cardiomyocytes was done using NIS Element software. The percentage of TUNEL-positive cardiomyocytes was calculated by quantifying the total number of cardiomyocytes present in a 20X area. All imaging and quantification was done in a blinded manner.

### Cell lysis preparation and Immunoblotting

Cell lysis preparation and immunoblotting was performed following the protocol described previously [8]. Briefly, post-treatment cells were quickly washed with ice-cold PBS and cell lysis buffer (Cell Signaling #9803), containing protease and phosphatase inhibitors cocktail (Sigma #P8340, #P0044), was added. Cells were scraped and collected in a tube followed by 3X vortexing every 5 minutes. Cell lysates were centrifuged at 15,000 *g* for 15 minutes at 4^0^C and supernatant was separated and, protein quantification was done using Bradford protein assay (Bio-Rad #5000001). SDS-PAGE was loaded with an equal amount of protein and was trans-blotted onto nitrocellulose membranes followed by incubation with primary antibody for overnight at 1:1,000 dilution for anti-p38α (Cell Signaling# 9218), phospho-p38 (Cell Signaling #9211), phospho-ERK1/2 (Cell Signaling #9101), ERK1/2 (Cell Signaling #4695), phospho-JNK (Cell Signaling #4668), JNK (Cell Signaling #3708) and 1:10,000 for GAPDH (Fitzgerald #10R-G109a). HRP labelled secondary antibodies Goat anti-Rabbit (Cell Signaling #7074) and Horse anti-Mouse (Cell Signaling #7076) were used at 1:3,000 dilution for 1 hour at room temperature. Membranes were developed using enhanced chemiluminescence (Thermo Scientific #32106) and scanned under the BioRad image analyzer.

### AC16 cell culture, transfection, and treatment

AC16 cardiomyocyte cell line derived from human ventricle was cultured in a DMEM/ Ham’s F-12 nutrient mixture (Sigma #D6434) containing 10% fetal bovine serum (FBS), 2mM L-glutamine and 1% penicillin/ streptomycin antibiotic cocktail (Sigma-Aldrich #P4333). The culture flasks were incubated in a 37°C CO_2_ humidified incubator containing 5% CO2 and 95% air. Cells were splitted at 80-90% confluency using 1X Trypsin (Sigma-Aldrich #T4299). Prior to transfection with either GFP containing control plasmid (Addgene #11153) or human *NRK2* containing plasmid (Addgene #23393), AC16 cardiomyocytes were serum-starved with 1X Opti-MEM (Gibco #00448) for 3 hours. Plasmid transfection was done using Fugene 6 (Promega #E2693) in a 3:1 ratio and incubated in CO_2_ incubator for another 3 hours. After 3 hours of transfection, complete DMEM/ Ham’s F-12 containing 2X FBS, penicillin-streptomycin and L-glutamine was supplemented and incubated for 24 hours in the CO_2_ incubator. Post-transfection, AC16 cells were treated with Ang II or vehicle control for 48 hours (for cardiac hypertrophy and RNA isolation) and 10 minutes for cell signaling studies. The experiment was repeated in duplicate using cells from two different passages.

### Wheat germ-agglutinin staining

Wheat germ agglutinin (WGA) staining was performed for cardiomyocyte circumference measurement as described previously [28]. Briefly, after deparaffinization, cardiac tissue (n=3 each sham group, n=6 each TAC group) rehydrated through serial concentration of ethanol. Antigen retrieval was performed using acetate-based retrieval solution followed by blocking the heart sections using animal-free blocking buffer (Cell signaling #15019). Sections were incubated with WGA-Alexa Fluor 549 (5μg/mL) dye for 15 minutes at room temperature followed by 3X washing with PBS and mounting using mounting media.

To analyze the role of NRK-2 in Ang II-induced hypertrophy in AC16 cardiomyocytes, an equal number of cells were seeded on the coverslip and transfected with NRK-2 or control plasmids as described above. Post-transfection, cells were treated with 1.0 μM Ang II or vehicle for 48 hours. After treatment, coverslips were washed with PBS and stained with WGA for 15 minutes at room temperature. Imaging was done using Olympus IX53 Inverted fluorescence microscope with a 40X object. Representative images were acquired under Nikon Eclipse Ti confocal microscope. Cell circumference was measured using NIS Element software. The experiment was repeated twice, in duplicate and triplicate using cells from two different passages.

### RNA isolation and quantitative real-time PCR (qRT-PCR)

RNA isolation from LV tissue and AC16 cardiomyocyte lysates was performed using an RNeasy kit (Qiagen #74104) following the instruction manual. Complementary DNA (cDNA) was synthesized using iScript™ cDNA Synthesis kit (BioRad #170-8891) following user’s guide and *NRK1, NRK2, atrial natriuretic peptide (ANP) and brain natriuretic peptide (BNP)* gene expression levels were assessed through qRT-PCR using Taqman Universal PCR Master Mix and gene expression assays (ABI, Carlsbad, CA) for NRK-1 (ID: Hs00944470_m1, Mm00521051_m1), NRK-2 (ID: Hs01043681_m1, Mm01172899_g1), ANP (ID: Mm01255748_g1, Hs00383230_g1) and BNP (ID: Mm00435304_g, Hs00173590_m1). The 18S rRNA assay probe (ABI #4319413E) was included in each reaction as endogenous control and to calculate the relative gene expression by using δ-δ Ct method.

### Annexin-V and 7-AAD labeling

An equal number of AC16 cardiomyocytes cells were seeded in 6-well plates and incubated in 37°C 5% CO_2_ humidified incubator. After transfection with control or NRK-2 plasmid, AC16 cardiomyocytes were serum-starved overnight followed by treatment with vehicle or 1μM Ang II. Post-48 hours of treatment, cells were trypsinized and harvested and pelleted by centrifugation. A total of 0.3 × 10^6^ cells were re-suspended in a fresh FACS tube using 100 µl of annexin binding buffer (Biolegend #B266205) containing annexin-V (Biolegend 640906) and 7-amino-actinomycin D (7-AAD) reagent (Biolegend 420403) and incubated for 30 minutes at room temperature in the dark. After incubation, 100 µl of 1X PBS containing 0.5 % BSA was added to each tube and vortexed quickly. Results were acquired using BD FACS Aria III Flow Cytometer and data was analyzed using FlowJo. The experiment was repeated thrice in duplicate using cells from three different passages.

### Mitochondrial membrane potential assay

The mitochondrial membrane potential (MMP) assay was carried out to assess the mitochondrial function, using the tetramethyl-rhodamine ethyl ester (TMRE) based MMP kit (Abcam, #ab113852) following the manufacturer’s protocol. Briefly, 0.3 × 10^6^ of AC16 cardiomyocytes were seeded in 6-well plates and incubated in 37°C, 5% CO_2_ humidified incubator. After transfection with control or NRK-2 plasmids, cells were serum-starved overnight followed by treatment with vehicle or 1 μM Ang II for 24h. The cells in a positive control group were then treated with 20 μM FCCP for 10 min which depolarizes the mitochondria. All the cells, except the DMSO group, were then treated with TMRE (1 μM) for 30 min and cells were rinsed with ice-cold 1X PBS in 0.2% BSA thrice. Cells were then harvested, pelleted, and washed with ice-cold 1X PBS containing 0.2% BSA. Finally, the cells were resuspended in 300 μl of ice-cold 1X PBS containing 0.2% BSA and subjected to Flow cytometry (BD FACS Aria III Flow Cytometer). Data analysis was done using FlowJo software.

### Statistics

Data group differences were evaluated for significance using one-way or two-way ANOVA followed by Tukey’s post-hoc test for multiple comparisons (Graph Pad Prism Software Inc., San Diego, CA). The specific statistical methods used are mentioned in each figure legend. Data are expressed as mean ± SEM. For all tests, a *P* value <0.05 was considered for statistical significance.

## Results

### Induction of NRK-2 but not NRK-1 in the pressure overloaded hearts

NRK-2 is predominantly expressed in skeletal muscle and trace amounts have been detected in healthy cardiac muscle [6, 8]. The expression of NRK-2 dramatically increases and coincides with HF following ischemia [8] or DCM [7, 29, 30]. Therefore, we hypothesized if cardiac pathogenesis dysregulates the expression of not only NRK-2 but both isoforms NRK-1 as well as NRK-2. To address this, a cohort of C57BL/6 mice was recruited and PO-model was generated through TAC surgery. Sham surgery was performed in an experimental control group. Post-3 weeks of TAC or Sham surgeries, expression levels of NRK-1 and NRK-2 was assessed in LV tissues. As expected, NRK-2 expression level was ∼5 fold higher in post-TAC vs. Sham hearts; however, interestingly, the expression level of NRK-1 was unchanged between Sham and post-TAC hearts (**Fig. 1A**). These results suggest that NRK-2 but not NRK-1 may have potential roles in PO-induced adverse cardiac remodeling and HF.

**Figure 1.**
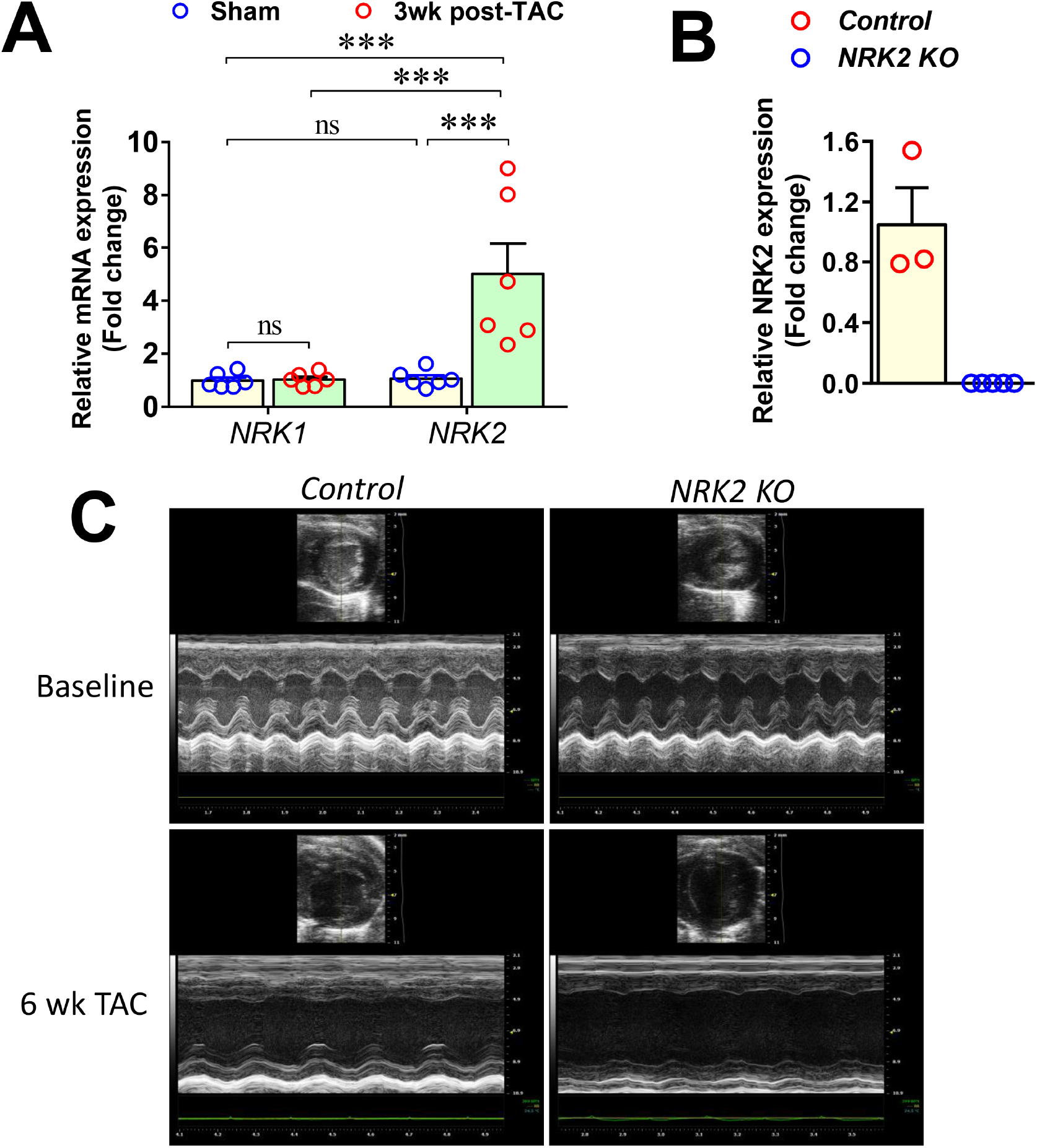
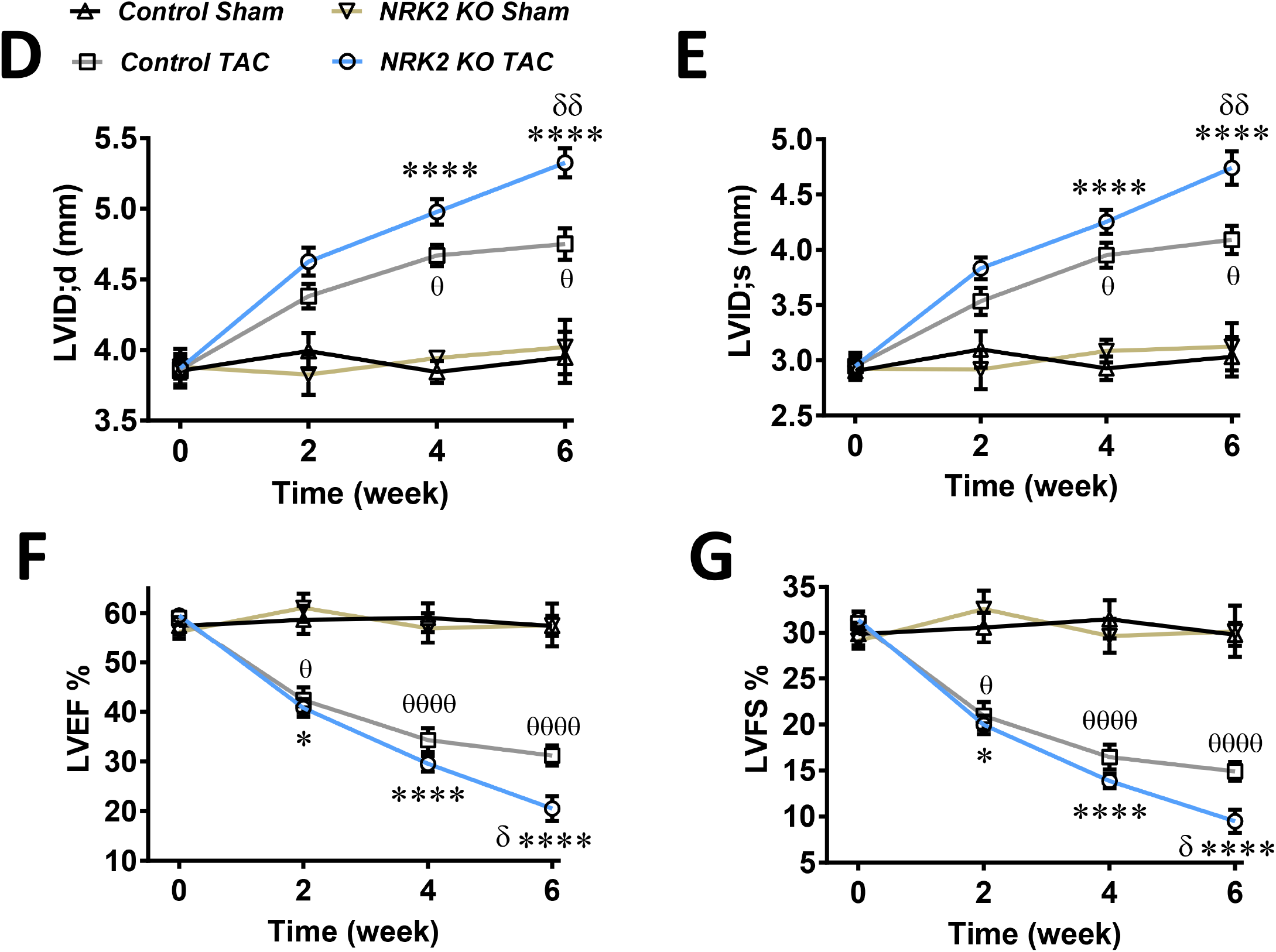
Loss of *NRK2* aggravates adverse cardiac remodeling and dysfunction post-TAC. (**A**) Bar diagram shows the relative expression of NRK-1 in Sham and trans-aortic constriction (TAC)-operated control hearts. The NRK-2 expression was significantly elevated in the left ventricle (LV) of mice subjected to TAC vs. Sham. (**B**) The *NRK2* mRNA expression was assessed in control and KO hearts and the bar diagram shows undetectable *NRK2* transcripts in the KO animals. (**C**) Representative images of B-mode and M-mode echocardiography from control and KO mice at baseline and 6 weeks post-TAC. Post-TAC echocardiography at 0, 2, 4 and 6 weeks shows (**D**) LV end-diastolic dimension (LVID;d) were comparable at 0 and 2 weeks and was significantly greater at 4 and 6 weeks in the NRK-2 KO animals post-TAC. (**E**) LVID;s was comparable between control and NRK-2 KO mice until 4 weeks and significantly dilated in the KOs at 6 week post-TAC. (**F**) LV ejection fraction (LVEF) and (**G**) LV fractional shortening (LVFS) were comparable until 4 weeks post-TAC and markedly deteriorated in the KO animals at 6 week post-TAC. Each dot of the scatter plots shows a mouse. A two-way ANOVA followed by Tukey’s multiple comparisons was performed to compare the group. δ, comparison between control-TAC vs. NRK-2 KO-TAC; *, comparison between Control-Sham vs. NRK-2 KO-TAC; θ, comparison between Control-Sham vs. Control-TAC.

### NRK-2 knockout exacerbates pressure overload -induced cardiac remodeling and dysfunction

We next asked whether NRK-2 overexpression in HF is compensatory or pathological. To address the same NRK-2 KO mouse was employed. First, the *NRK2* gene expression level in the KO and littermate controls was assessed which confirms the undetectable *NRK2* mRNA transcripts in the KO animals (**Fig. 1B**). A batch of *NRK2* KO and littermate control mice was recruited and TAC or Sham surgeries were performed. Cardiac function was assessed through trans-thoracic M-mode echocardiography using Vevo2100 at 0-, 2-, 4- and 6-weeks post-TAC (**Fig. 1C**). The reduction in LV ejection fractions (LVEF) and LV dilatation in NRK-2 mice was similar to *wild-type* mice for the first two weeks. However, the NRK-2 mice showed an accelerated decline in LVEF and LV fractional shortening (LVFS) along with an expansion of diastolic LV interior dimension (LVID;d) for the subsequent two weeks, which were further worsened during weeks five and six post-TAC (**Fig. 1D-G**). Overall post-TAC mortality was comparable in both groups (**Suppl. Fig. 1**). These data suggest that the protective effects of NRK-2 are minimal in the early hypertrophic remodeling phase of PO, but is protective against chronic PO-induced LV dilatation and cardiac dysfunction.

### Loss of NRK-2 aggravates pressure overload-induced cardiac hypertrophy and heart failure

To identify the specific roles of NRK-2 in PO-induced cardiac remodeling and heart failure, heart and lung tissues were evaluated morphometrically at six-week post-TAC. Heart weight (HW) and lung weight (LW) to tibia length (TL) ratios revealed comparable values in the sham-operated mice. As expected, HW/TL ratio was higher in TAC-operated animal groups in comparison to the sham group, however, the ratio was significantly lower in the control mice vs. *NRK2* KO mice subjected to TAC (**Fig. 2A**). Moreover, LW/TL ratio was significantly higher in the KO compared to control mice subjected to TAC though the ratio in the control mice was comparable to the sham-operated group which indicates that control mice subjected to TAC did not develop PO-induced HF at six week (**Fig. 2B**). The heart sizes were smaller in the sham-operated vs. TAC groups. In the sham-operated group, we did not find a difference between the control and NRK-2 KO mice. In the TAC group, the heart sizes of NRK-2 KO mice were significantly larger than the control mice (**Fig. 2C**). This was associated with dilated LV chambers in the KO vs. control hearts at six-week post-TAC (**Fig. 2D**). At the cellular levels, the cardiomyocytes of sham-operated control and NRK-2 KO mice were similar in size and smaller than the TAC group (**Suppl Fig. 2A**). Among the TAC mice, cardiomyocytes from NRK-2 KO mice showed significantly more hypertrophy than the control group (**Fig. 2E-F**). Next, we assessed the molecular markers of cardiac hypertrophy, including atrial natriuretic peptide (ANP) and brain natriuretic peptide (BNP). As expected, irrespective of genotype, the expression of both peptides was upregulated in the mice subjected to TAC vs. sham group. However, the expression of ANP and BNP were higher in the NRK-2 KO vs. control hearts subjected to TAC (**Fig. 2G-H**). Taken together, these findings suggest that NRK-2 protects the heart against chronic PO-induced cardiac hypertrophy and heart failure.

**Figure 2.**
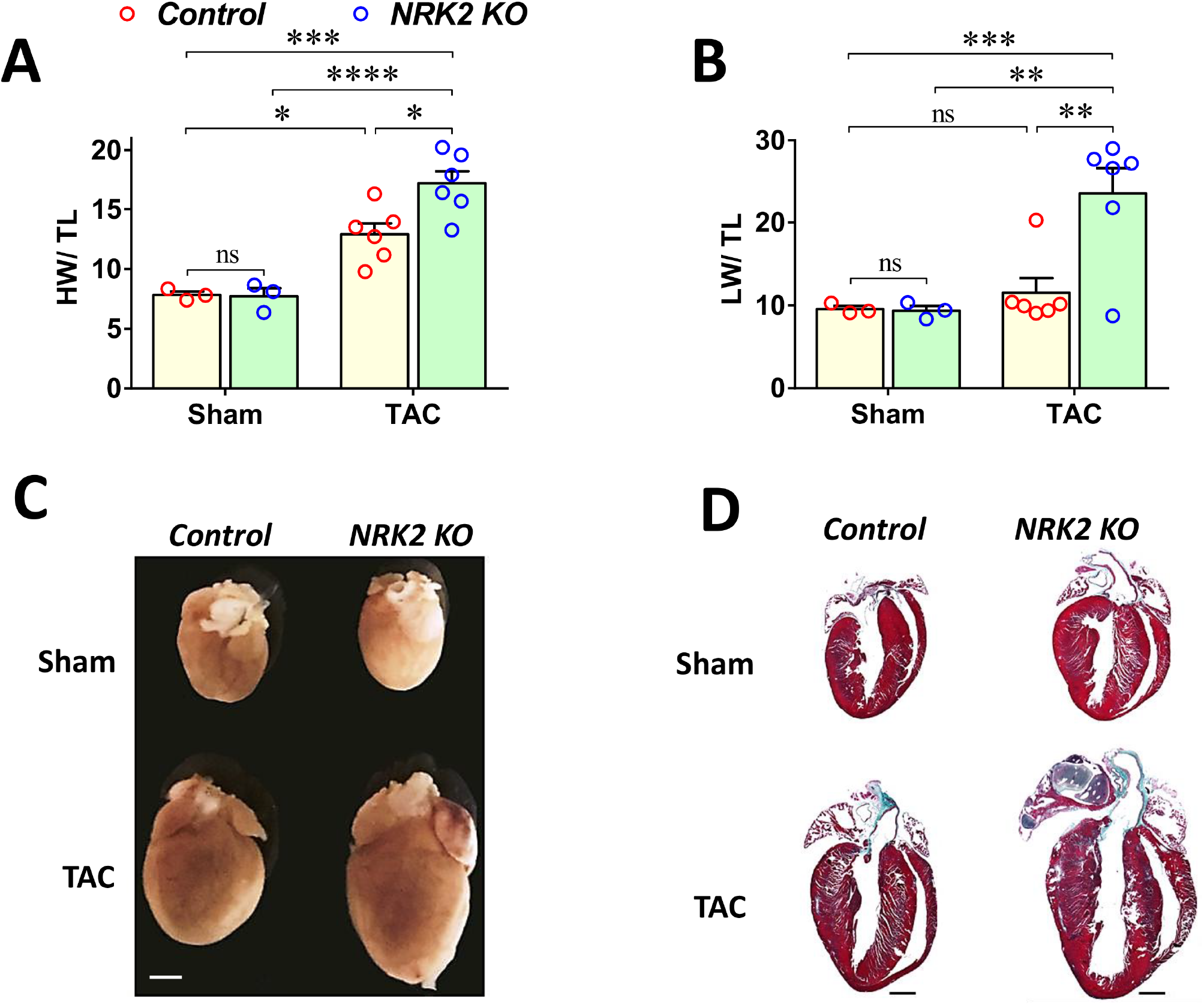
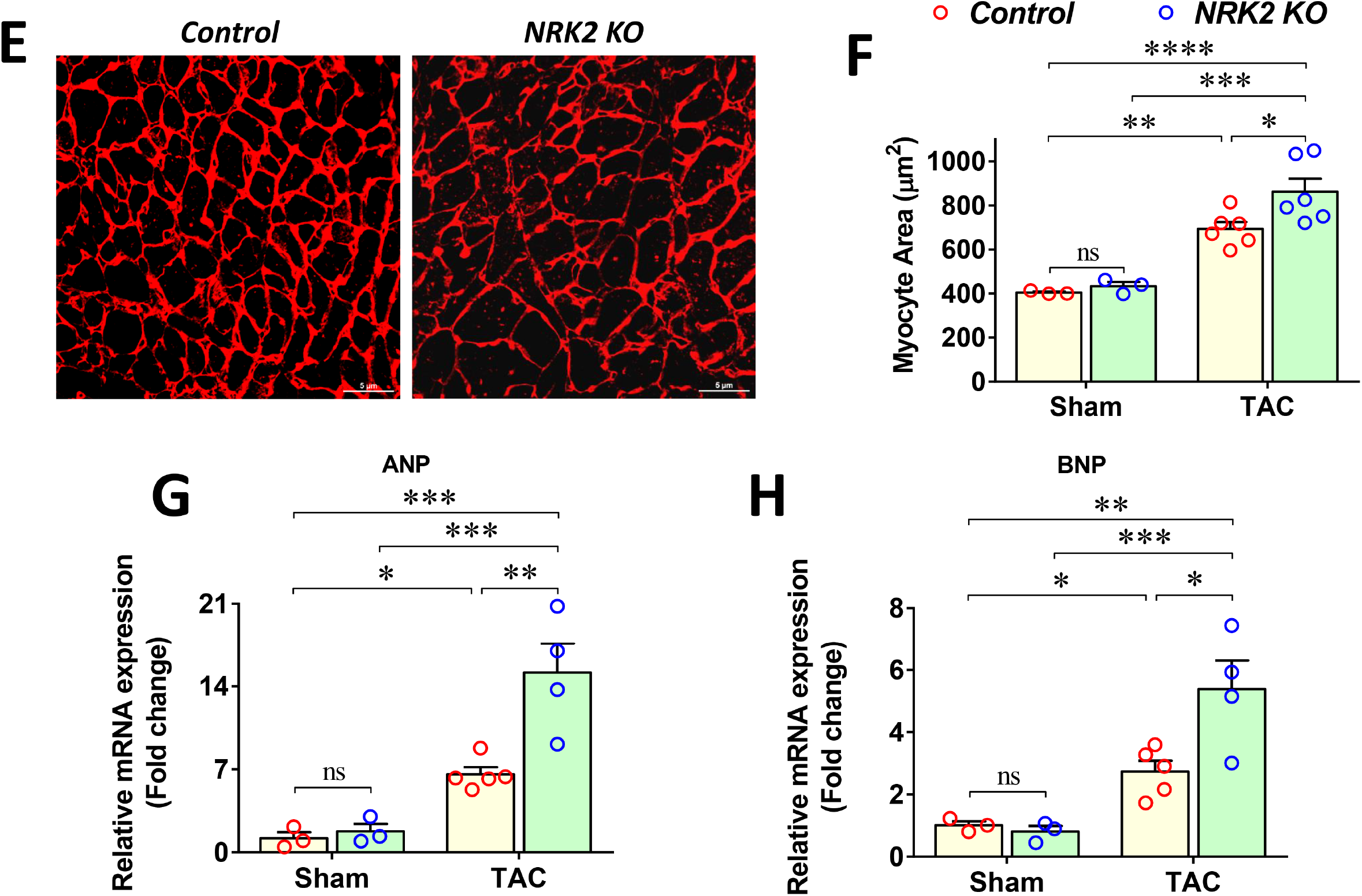
NRK-2 protects against TAC-induced cardiac hypertrophy, LV chamber dilatation and heart failure. Morphometric analysis reveals a significantly increased (**A**) heart weight (HW) to tibia length (TL) and (**B**) lung weight (LW) to tibia length ratios in the NRK-2 KO vs. control mice post-6 weeks post-TAC. (**C**) Representative formalin-fixed control and KO heart images from sham and 6 weeks post-TAC groups. Scale bar, 1mm. (**D**) Representative Masson’s trichrome-stained longitudinal heart sections show comparable LV chambers in Sham-operated control and KO groups, however, marked LV chamber dilatation in the KO vs. control heart post-TAC. Scale bar, 1mm. (**E**) Representative images of wheat germ agglutinin (WGA) stained heart sections from control and KO mice subjected to TAC. Scale bar, 5μm. (**F**) The cross-sectional areas of cardiomyocytes were comparable in the sham-operated control and NRK-2 mice but significantly increased in the NRK-2 KO *vs*. control group post-TAC. (**G**) Bar diagram shows an increased expression level of fetal genes atrial natriuretic peptide (ANP) and (**H**) brain natriuretic peptides (BNP) in the KO vs. control hearts post-TAC. Each dot of the scatter plots shows a mouse. A two-way ANOVA followed by Tukey’s multiple comparisons was performed to compare the group. ns; not significant; *p<0.05; **p<0.005; ***p<0.005; ****p<0.0001.

### Genetic ablation of NRK2 exacerbates dilated cardiomyopathy and myocardial fibrosis post-TAC

Chronic PO eventually leads to cellular apoptosis which induces fibroblast activation and extracellular matrix deposition in the heart. To evaluate the impact of NRK-2 deletion on cardiomyopathy and fibrotic cardiac remodeling post-TAC, we performed TUNEL staining using 6 weeks post-TAC NRK-2 KO and control mouse heart. Analysis showed a comparable number of TUNEL-positive cardiomyocytes in sham-operated groups. As expected, irrespective of genotype the number of TUNEL-positive cardiomyocytes was significantly increased in the TAC vs. sham groups. However, the percentage of apoptotic cardiomyocytes in the KO animals was profoundly higher in comparison to the control groups subjected to TAC (**Fig. 3A-B, Suppl Fig. 2B**)). Since fibrotic tissues eventually replace the apoptotic cells, we next assessed cardiac fibrosis using Masson’s trichrome staining. Consistent with cardiomyocyte apoptosis, a comparable fibrotic area was observed between the sham-operated groups, and it was significantly increased in both TAC-vs. sham-operated groups. Interestingly, the fibrotic area was markedly greater in the KO compared to control hearts following TAC (**Fig. 3C-D, Suppl Fig. 2C**). The observed data strongly suggest that NRK-2 protects the heart from dilatative cardiomyopathy and its pathological consequences including cellular apoptosis and cardiac fibrosis.

**Figure 3.**
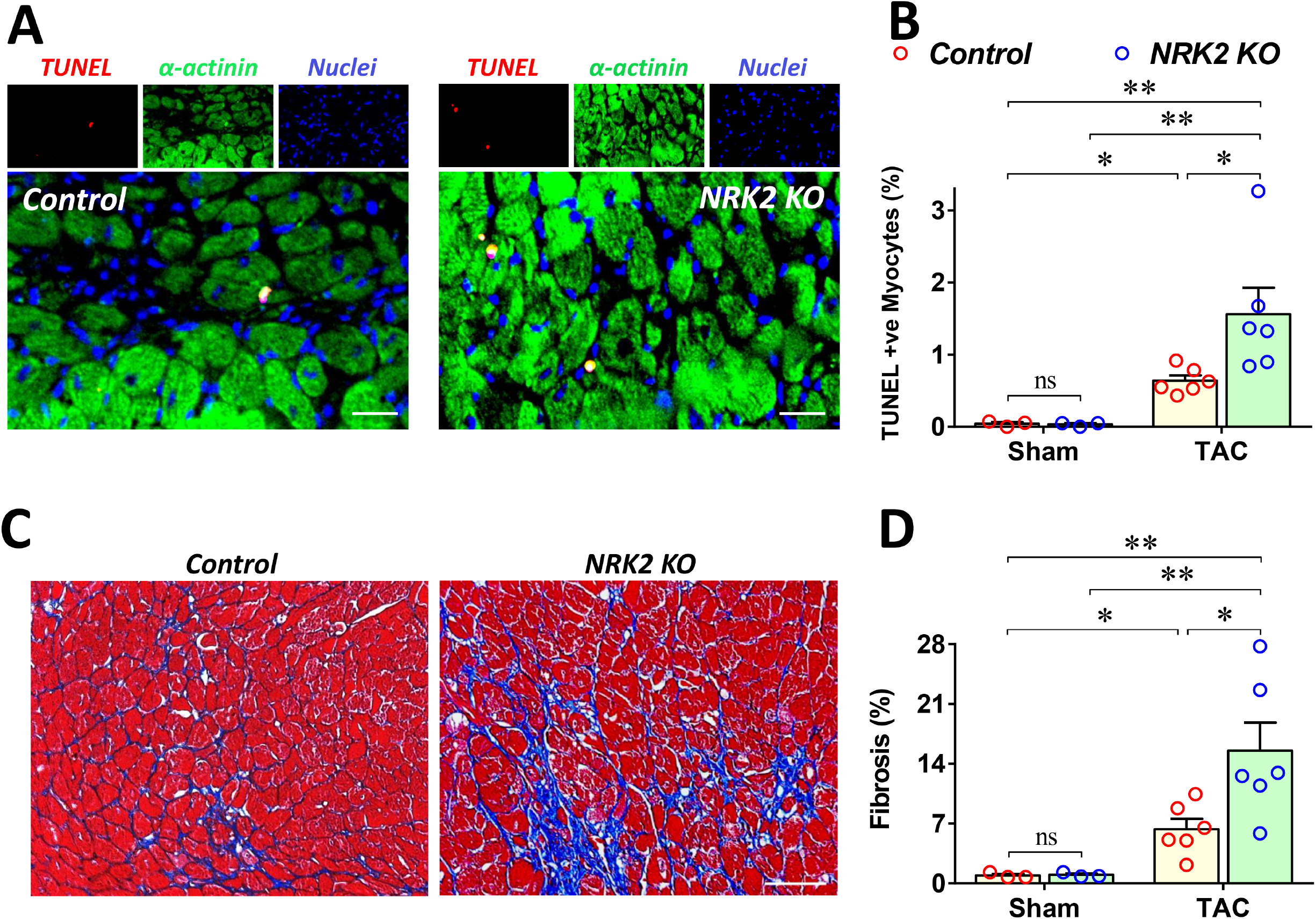
Loss of NRK-2 exacerbates dilated cardiomyopathy and cardiac fibrosis post-TAC. **(A)** Representative images of TUNEL positive (apoptotic) cardiomyocytes from KO and littermate control hearts post-TAC. Scale bar, 20μm. **(B)** Quantification shows a significantly increased proportion of apoptotic cardiomyocytes in the NRK-2 KO mice in comparison to the control group, n=7 each, *P<0.05. **(C)** Representative images of Masson’s trichrome-stained LV from 6 weeks post-TAC NRK-2 KO and control hearts. Scale bar, 60μm. **(D)** Quantification of the fibrotic area shows a significantly higher percentage of fibrotic tissues in the NRK-2 KO *vs*. control hearts post-TAC. Each dot of the scatter plots shows a mouse. A two-way ANOVA followed by Tukey’s multiple comparisons was performed to compare the group. *P<0.05; **p<0.005.

### NRK-2 overexpression attenuates angiotensin II-induced cardiomyocyte hypertrophy and fetal gene expression

Based on findings that NRK-2 deficiency promotes cardiac hypertrophy and dilated cardiomyopathy post-TAC, we hypothesized that NRK-2 gain-of-function attenuates Ang II-induced cardiomyocyte hypertrophy. In this series, AC16 cardiomyocytes were first transfected with human *wild-type NRK2* or *GFP* plasmids and mRNA expression level was assessed. NRK2 expression was profoundly upregulated in the AC16 cells transfected with *NRK2* plasmids vs. the *GFP* group. (**Fig. 4A**). Next, transfected AC16 cells seeded on a coverslip were treated with Ang II followed by WGA staining and cell circumference was measured. Analysis revealed that cardiomyocyte sizes were comparable between the control and NRK-2 overexpressing group, however, consistent with *in vivo* findings, cardiomyocytes with NRK-2 overexpression were significantly smaller than the control group treated with Ang II (**Fig. 4B-C**). These observations further attest that NRK-2 critically protects against the PO-induced cardiac hypertrophy.

**Figure 4.**
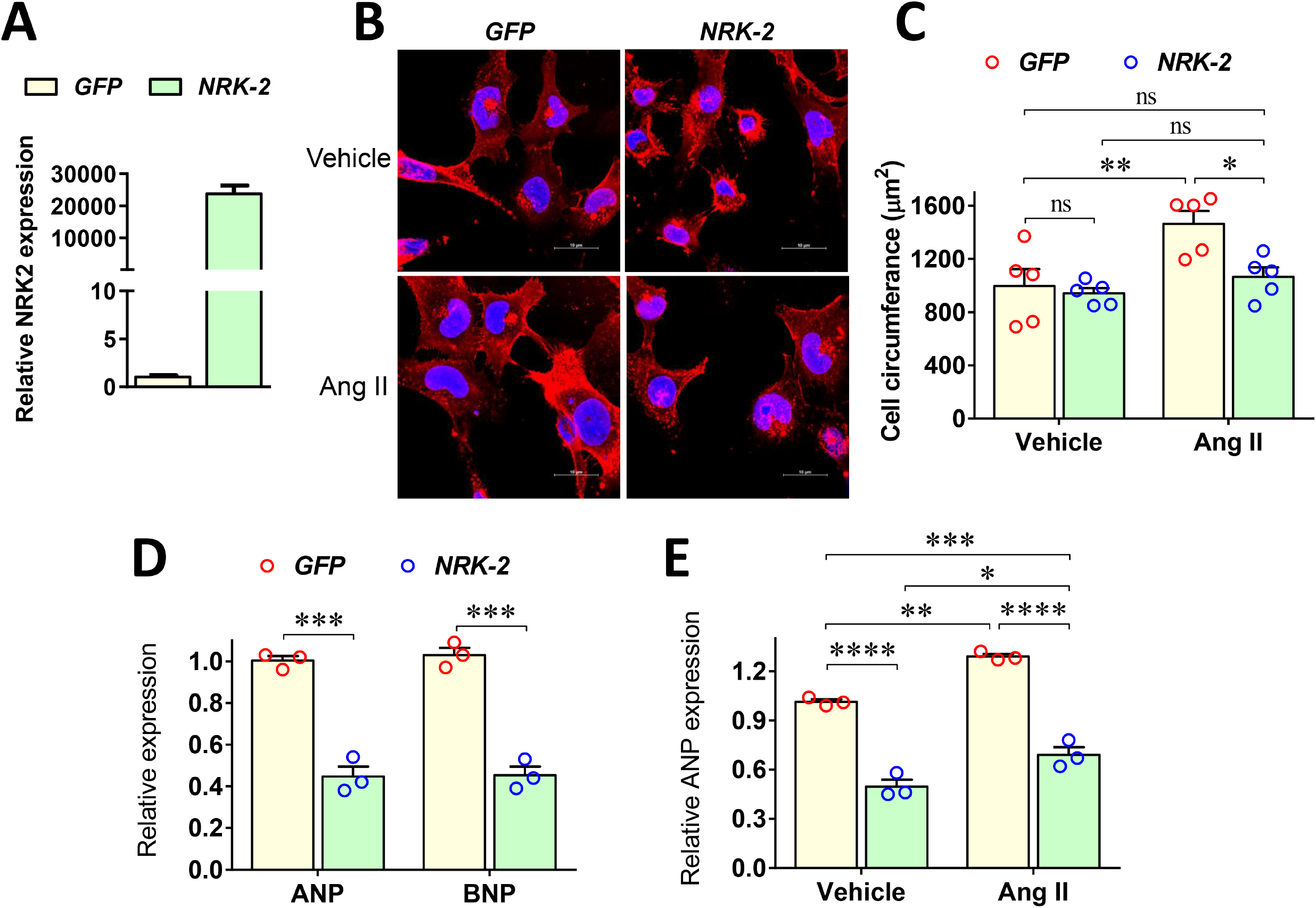
Gain-of-NRK-2 function attenuates angiotensin II- induced cardiomyocyte hypertrophy and suppresses fetal genes expression. (**A**) The graph shows the level of NRK2 expression in GFP or NRK2 overexpressing AC16 cardiomyocytes. (**B**) Representative images show wheat germ agglutinin (WGA)-stained AC16 cardiomyocyte. Scale bar, 10μm (**C**) Cross-sectional areas of cardiomyocyte show a significantly smaller cell size in the NRK-2 overexpressing group vs. control treated with Angiotensin II (Ang II) for 48 hours. **(D)** The bar diagram shows significantly attenuated atrial (ANP) and brain natriuretic peptides (BNP) expression in the NRK-2 overexpressing AC16 cardiomyocytes. An unpaired *t*-test was performed to compare the group. **(E)** The bar diagram shows that Ang II treatment further increases the ANP expression in cardiomyocytes which was significantly abrogated in NRK-2 overexpressing cells. Each dot of the scatter plots shows replicates of the experiments. A two-way ANOVA followed by Tukey’s multiple comparisons was performed to compare the group. *P<0.05; **P<0.005; ***p<0.005; ****p<0.0001.

Fetal gene expression correlates with heart failure in a variety of cardiac pathogenesis [31, 32]. Consistently, we observed that *NRK2* KO hearts displayed induction of fetal genes *ANP* and *BNP* expression post-TAC (**Fig. 2G-H**). We consequently hypothesized that NRK-2 overexpression in AC16 cardiomyocytes suppresses *ANP* and *BNP* genes expression. AC16 cells were transfected with *NRK2* or control plasmids followed by measurements of *ANP* and *BNP* mRNA expression. Consistent with our *in vivo* data, overexpression of NRK-2 in the cardiomyocytes significantly downregulated *ANP* and *BNP* expressions levels (**Fig. 4D**). Next, we tested if NRK-2 overexpression limits *ANP* expression in a pathological hypertrophy model. In this series, NRK-2 overexpressing and control AC16 cells were treated with Ang II, and the *ANP* mRNA expression was assessed. The expression of *ANP* has further increased in both cell groups post-Ang II treatment however, the pattern was similar to as seen between the vehicle-treated groups (**Fig. 4E**). These findings suggest that NRK-2 not only protects against pathological cardiac hypertrophy but also limits heart failure.

### NRK-2 gain-of-function attenuates angiotensin II-induced mitochondrial membrane depolarization and cardiomyocyte death

It is well established that the renin-angiotensin system plays a crucial role in the development of PO– induced hypertrophy and hypertrophic cardiomyopathy. Therefore, we next asked whether the NRK-2-induced down-regulation of HF markers is associated with reduced cell death in the AC16 cells. Consistent with the results from the loss-of-function *in vivo* model, NRK-2 overexpression protected from Ang II-induced cell death. The cell viability was found to be comparable between vehicle-treated NRK-2 and control groups; however, viability was markedly higher in the NRK-2 overexpressing cardiomyocyte vs. control cells treated with Ang II (**Fig. 5A-B**). In contrast, the relative proportion of ann-V-(apoptotic) or 7-AAD-positive (necrotic) cells was markedly lower following the NRK-2 overexpression in comparison to the control group treated with Ang II (**Fig. 5C-D**). Similarly, the percent of dual-positive (late apoptotic) cells was significantly lower in the NRK-2 overexpression group (**Fig. 5E**).

**Figure 5.**
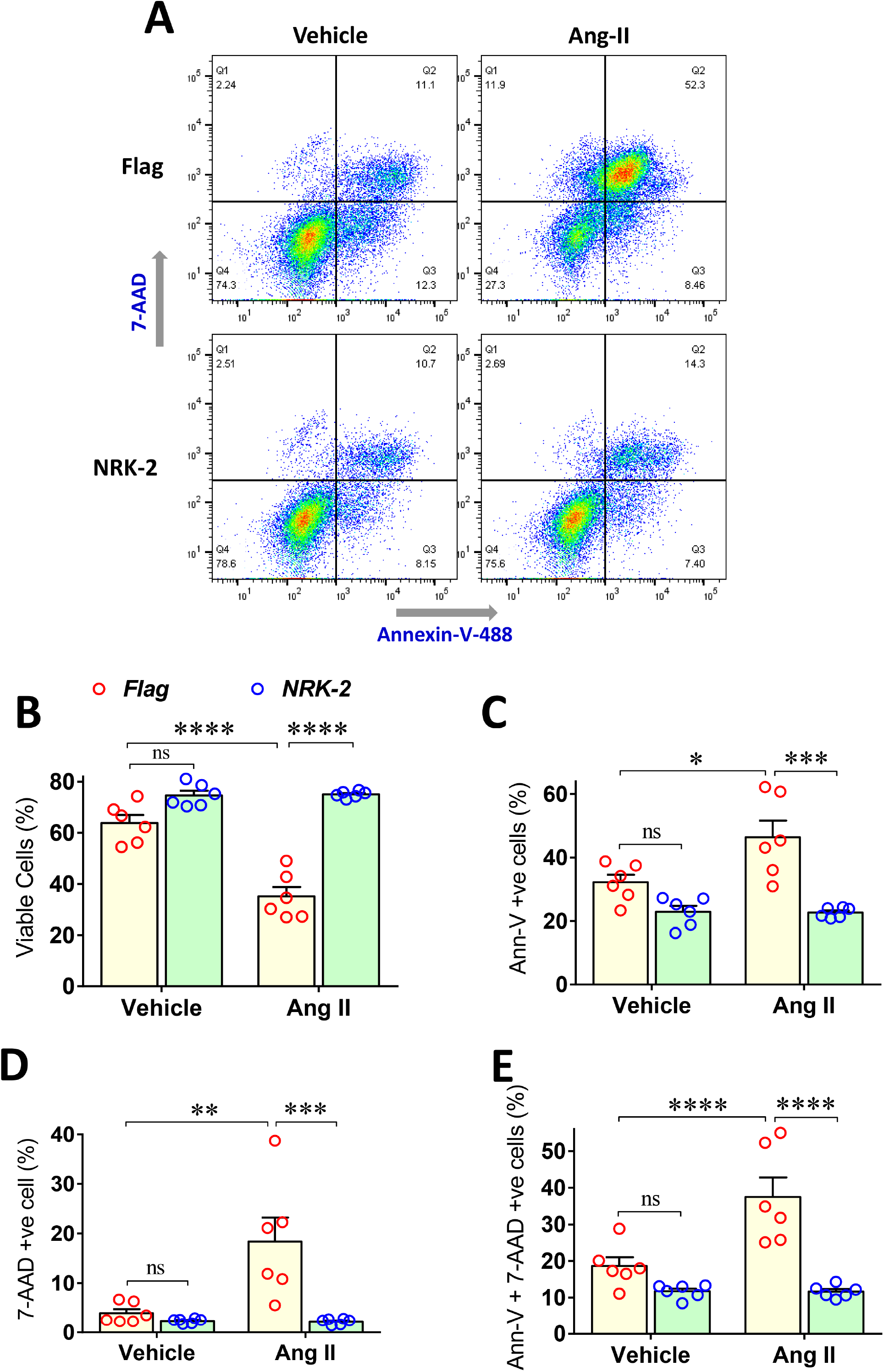
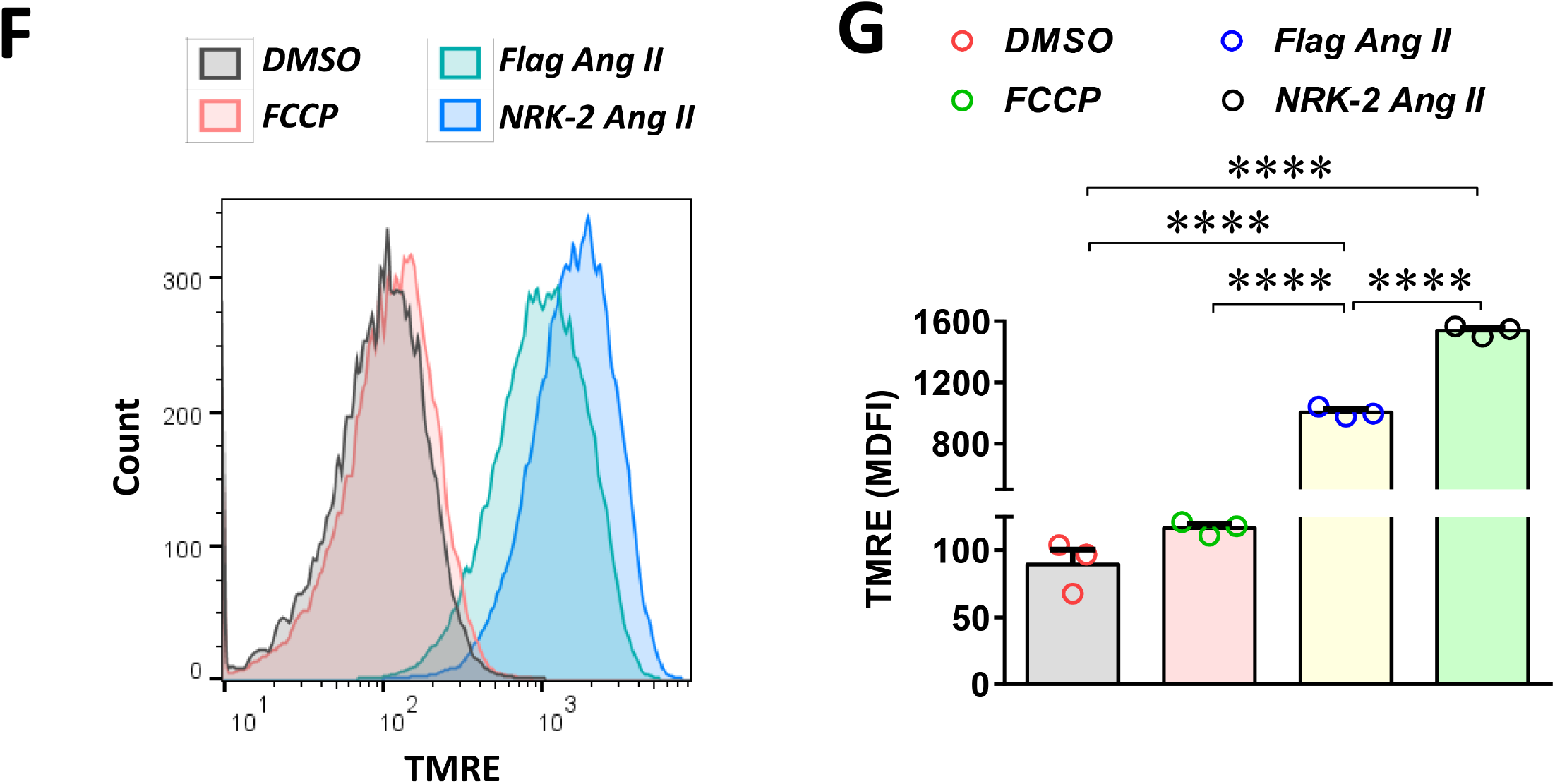
NRK-2 protects against angiotensin II-induced AC16 cardiomyocytes death. (**A**) Representative flow cytometry scatter plots show annexin V-Alexa-488 and 7-AAD positive cardiomyocytes in Flag control and NRK-2 overexpressing groups challenged with angiotensin II (Ang II). (**B**) AC16 cells were transfected with NRK-2 or control plasmids and challenged with Ang II for 48 hours. Flow cytometry analysis shows that NRK-2 overexpression significantly increased the cell viability post-Ang II treatment. Further analysis shows a significantly decreased level of (**C**) annexin-V-positive (apoptotic), (**D**) 7-AAD-positive (necrotic), and (**E**) dual-positive (late apoptotic) cells in NRK-2 overexpression vs. control group. (**F**) Representative histograms show tetramethyl-rhodamine ethyl ester (TMRE) fluorescence intensity in DMSO and FCCP controls and, Ang II treated groups (**G**) Bar diagram from median fluorescence intensity (MDFI) shows a significantly better mitochondrial membrane potential (MMP) or mitochondrial function in the NRK-2 overexpressing cells vs. control group treated with Ang II. Each dot of the scatter plots shows replicates of the experiments. A two-way ANOVA (Ann-V experiment) and one-way ANOVA (TMRE experiment) followed by Tukey’s multiple comparisons were performed to compare the group. *P<0.05; **P<0.005, ***P<0.0005, ****P<0.0001.

Loss of mitochondrial membrane potential is one of the key factors in determining the onset of cell death. The strong involvement of NRK-2 as a pro-survival protein may have a significant role in protecting mitochondria. Hence, we examined the role of NRK-2 in the regulation of mitochondrial function under cardiac pathological conditions. In this series, AC16 cells were transfected with control or NRK-2 plasmid and treated with Ang II. MMP, required for mitochondrial function, measurement revealed a significantly higher MMP (better mitochondrial function) in the cells overexpressing NRK-2 compared to control cells treated with Ang II-induced (**Fig. 5F-G**). These results strongly suggest that gain-of NRK-2 function limits Ang II-induced mitochondrial dysfunction and cardiomyocyte death.

### NRK-2 suppresses Ang II-induced JNK activation

In PO condition, integrins and integrin-binding proteins play crucial roles in maladaptive dilatative cardiac remodeling [33-35] through regulation of various cardiomyocyte signaling including Akt and MAPK pathways [33, 35, 36]. Therefore, we sought to identify the possible roles of NRK-2 in the regulation of these signaling pathways under pathological conditions by employing the gain-of-function model in AC16 cardiomyocytes. To test this, we first determined the peak of JNK activation in AC16 cardiomyocytes post-Ang II treatment. The cells were treated with Ang II for different time points and the phosphorylation level of JNK (Thr183/Tyr185) and P38 (Thr180/Tyr182) was assessed. The phosphorylation was rapidly increased and peaked at 10 minutes for JNK and 15 minutes for P38 post-Ang II treatment (**Fig. 6A-B**). Therefore, to assess the role of NRK-2 in the regulation of MAPKs, cells were transfected with GFP or NRK-2 and treated with Ang II for 10 minutes and phosphorylation level of MAPKs including JNK, P38 and ERK was assessed. NRK-2 overexpression in cardiomyocytes was found to have a minimal basal effect on JNK activity. However, NRK-2 gain-of-function significantly suppressed the JNK activation in cardiomyocytes post-Ang II treatment (**Fig. 6C-D**). Interestingly, the phosphorylation levels of P38 and ERK were higher in Ang II treated compared to vehicle-treated cells. However, the phosphorylation levels were comparable in control vs. NRK-2 overexpressing cardiomyocytes post-Ang II stimulation (**Fig. 6C-D**). These results strongly suggest that NRK-2 specifically regulates JNK MAPK in human cardiomyocytes in a model of PO. The gain-of-NRK-2 function profoundly inhibits JNK phosphorylation which potentially attenuates cardiomyocytes death post-Ang II –treatment.

**Figure 6.**
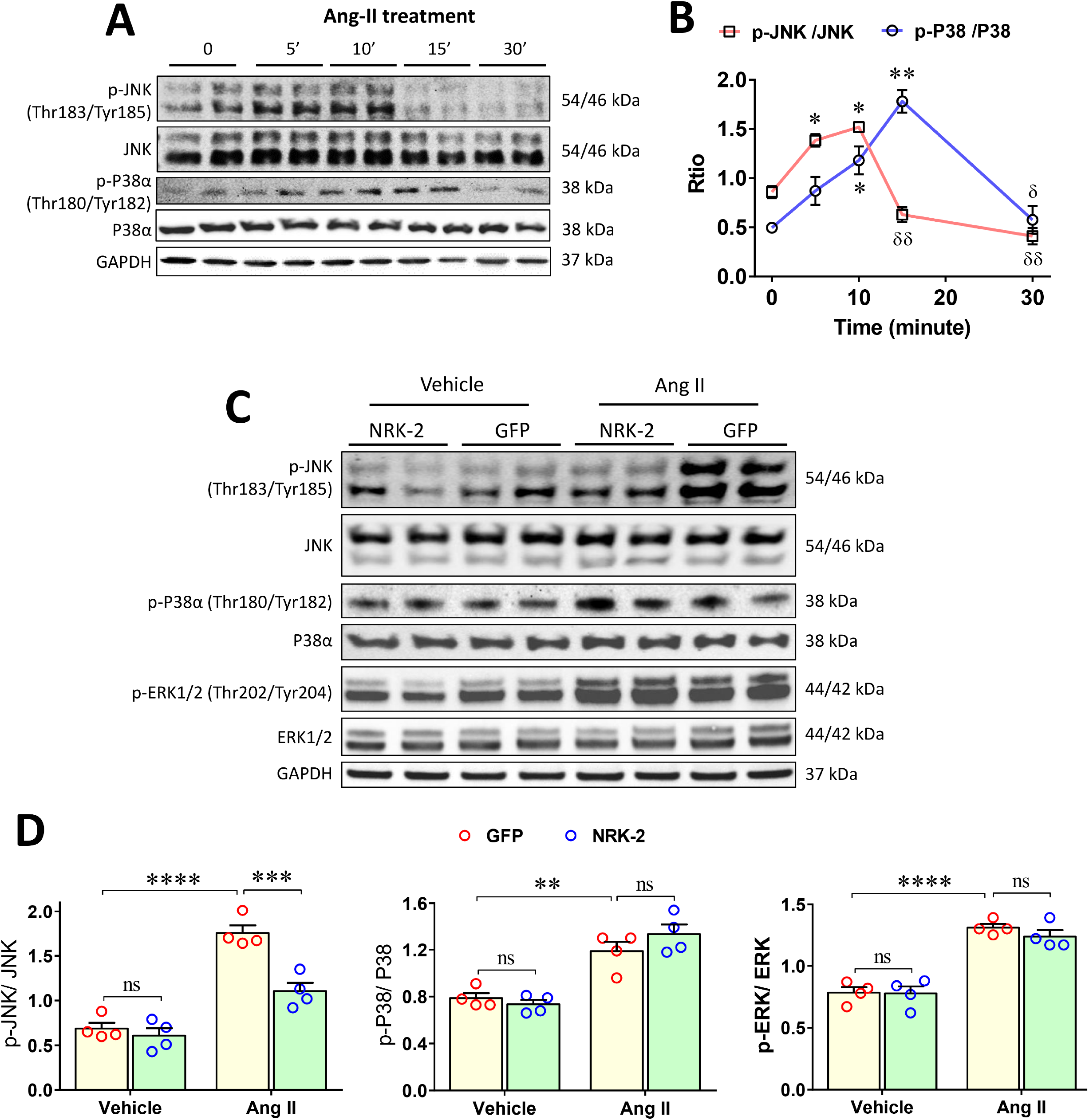
NRK-2 suppresses Ang II- induced JNK activation. **(A)** Representative blot images of p-JNK and p-P38α. (**B**) Blot quantification shows JNK and P38α phosphorylation (activation) in control cardiomyocytes treated with Ang II for mentioned time points. JNK phosphorylation rapidly increases and peaked at 10 minutes and P38α phosphorylation peaked at 15 minutes post-Ang II (1μM) treatment and was quickly abolished. *, comparison between 0 and peak time points; δ, comparison between peak and time points where phosphorylation abolished. (**C-D**) Representative blots and scatter dot plots show a comparable p-JNK level in vehicle-treated GFP and NRK-2 overexpressing cardiomyocytes and a significantly lower level in NRK-2 overexpressing cells vs. GFP groups treated with Ang II for 10 minutes. The activity of P38 and ERK in Ang II or vehicle-treated NRK-2 overexpressing and control cardiomyocytes were comparable. Each dot of the scatter plots shows replicates of the experiments. A two-way ANOVA followed by Tukey’s multiple comparisons was performed to compare the group. Ang II, angiotensin II; *P<0.05; **P<0.005.

## Discussion

DCM is one of the most prevalent causes of heart failure and the prognosis remains poor even with the most advanced therapeutic approaches. The underlying molecular mechanisms leading to the development of DCM remain largely undefined. Understanding the pathomechanisms of DCM and identifying the therapeutic targets is of great clinical importance. Owing to the critical role of integrins in coupling mechanical stress with mechano-transducing the signaling pathway, we investigated the potential role of NRK-2, an integrin-binding protein, in the regulation of chronic PO-induced cardiac remodeling and underlying dysregulated cardiac cell signaling. We observed, for the first time, that NRK-2 critically regulates JNK pathway in Ang II stimulated human cardiomyocytes. Employing NRK-2 KO mice, here we show a time-dependent deterioration of cardiac function post-TAC, which is reflected by LV chamber dilatation, cardiac hypertrophy, DCM, fibrosis, and induction of heart failure markers (**Fig. 7**).

**Figure 7.**
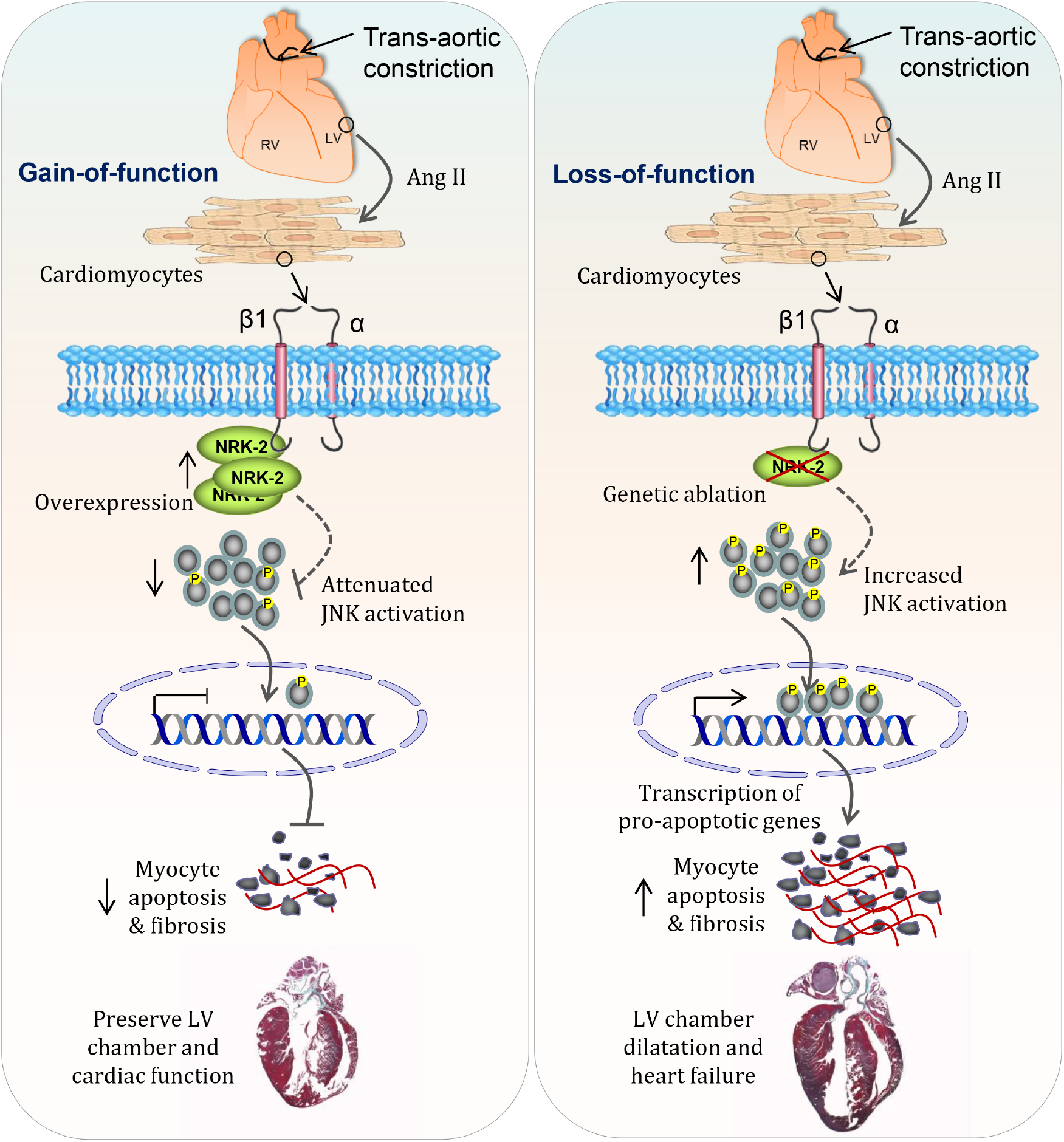
Schematic diagram shows the proposed molecular mechanism of NRK-2 mediated heart failure under pressure overload: Diagram, drawn based on our *in vivo* and *in vitro* findings, shows the role of NRK-2 in a pressure overload (PO) model–induced c-Jun N-terminal kinase (JNK) pathway modulation in the cardiomyocytes. PO induces overexpression of NRK-2 in the heart potentially inhibits the JNK pathway and limits dilatative cardiomyopathy. This was confirmed in an *in vitro* model employing gain-of-NRK-2 function in the human cardiomyocytes. NRK-2 upregulation attenuates JNK activation and cardiomyocyte death when treated with angiotensin II (Ang II). These findings suggest that loss of NRK-2 in the heart likely promotes JNK activation, dilatative cardiomyopathy, and heart failure. **β1**; β1 integrin, **α**; α integrin, **P**; phosphate group.

Recently, we reported that NRK-2 deficiency accelerated cardiac dysfunction following myocardial ischemia [8]. This profound detrimental cardiac phenotype was primarily associated with enhanced P38 MAPK activation. Interestingly, current findings from the PO model display a relatively delayed onset of adverse cardiac remodeling and dysfunction in the NRK-2 KO mice. In contrast to ischemic heart, Ang II treatment leads to JNK but not P38 or ERK inhibition in the NRK-2 overexpressing human cardiomyocytes. In complete agreement with our current findings, a recent report employing an acute PO model (2 weeks) has shown a trend towards LV chamber dilatation with comparable ERK activation in the NRK-2 KO mice. However, contrary to our findings, this study reported unchanged cardiomyocyte hypertrophy and ANP gene expression post-TAC [9]. These differential findings could be due to the differences in experimental settings as our studies mainly focused on the chronic maladaptive phase of cardiac remodeling post-TAC.

Multiple studies have shown the crucial roles of integrin and integrin-binding proteins such as Talin, Vinculin, Melusin, focal adhesion kinase (FAK) and integrin-linked kinase (ILK) in the pathogenesis of cardiac diseases through regulation of MAPKs [33, 35, 37-40]. Consistent with our *in vivo* and *in vitro* findings, mice with deficiency of melusin, another muscle-specific integrin-binding protein, displayed a similar detrimental cardiac phenotype when challenged with PO. Melusin deficiency leads to DCM and cardiac dysfunction post-TAC which at the molecular level was associated with an increased GSK-3β activation [40]. Consistently, *in vivo* gain-of-function model of melusin showed protective phenotypes where Melusin overexpression in the heart presented preserved LV chamber and contractile function under PO which found to be regulated through profound inhibition of GSK-3β and activation of ERK [34]. Later, a genetic study revealed that a point mutation in Melusin-encoding gene *ITGB1BP2* lead to DCM and severe impairment of left ventricular function [41].

Similar to NRK-2, the expression of Talin1 was upregulated in failing heart and no overt basal effect was seen in the cardiomyocyte-specific KO mice. However, unlike the phenotype observed in the NRK-2 KO mice, cardiomyocyte-specific loss of Talin1 attenuates cardiac hypertrophy, fibrosis and preserves cardiac function when challenged with PO. The observed phenotype from Talin1 KO was driven by the inhibition of ERK, P38 and Akt and activation of GSK-3β [42]. The reported detrimental phenotypes from loss-of-NRK-2 function mice models and other integrin-binding proteins are in complete agreement with the cardiac phenotypes observed in β1 integrin deficient mice. Alteration in β1 integrin function caused DCM and fibrosis associated with reduced cardiac contractility. Moreover, cardiomyocyte-specific β1 integrin KO mice demonstrated a poor survival rate post-TAC [43-45].

The adverse cardiac phenotypes in the mice without integrins or integrin-binding proteins are predominantly regulated by ERK and P38 MAPKs pathways. Interestingly, we observed that NRK-2 minimally regulates ERK and P38 activation in cardiomyocyte post-Ang II and suppression of JNK signaling emerged as the central molecular mechanism behind the cardioprotective effects of NRK-2. The role of JNK signaling pathways in the regulation of cardiac hypertrophy in response to PO has been the focus of many investigations. For instance, c-Jun (a downstream target of JNK) facilitates insulin-like growth factor (IGF)-Akt signaling leading to hypertrophy development and heart failure, while down-regulation of c-Jun signaling prevents cardiac hypertrophy [46]. Moreover, gain-of-MKK7 function (an upstream JNK-activator) in transgenic mice resulted in cardiac hypertrophy and heart failure, owning to excessive activation of JNK [47].

Our previous and current findings indicate that NRK-2 differentially regulates MAPK signaling in cardiac ischemia and PO conditions. The observations from the MI model revealed that P38 activation in the ischemic heart produced a severe detrimental phenotype and led to rapid heart failure post-MI. However, in the TAC model where detrimental cardiac phenotypes progressed gradually and JNK inhibition was seen in NRK-2 overexpressing cardiomyocytes post-Ang II treatment. These findings strongly suggest that NRK-2 regulates differential MAPK signaling in differential cardiac pathogenesis. It is also important to mention that P38 activation was observed in the ischemic NRK-2 KO mice, and JNK suppression was seen in NRK-2 overexpressing human cardiomyocytes post-Ang II treatment. Therefore, there is a possibility that NRK-2 may regulate P38 only in the mouse heart and JNK, particularly in human cardiac cells under pathological conditions.

Mitochondrial dysfunction plays a crucial role in cardiomyopathy under cardiac pathological conditions. The mitochondrial JNK activation is reported to trigger cellular apoptosis and exacerbates myocardial injury upon cardiac stressor [48]. Consistent with this finding, we identified that NRK-2 inhibits Ang II-induced JNK activation, mitochondrial dysfunction and cell death. Further investigation is needed to identify whether NRK-2 regulates only mitochondrial JNK, cytoplasmic JNK, or JNK located in both cellular compartments.

In conclusion, we demonstrated the critical roles of NRK-2 in chronic PO-induced dilated cardiac remodeling and heart failure. Employing NRK-2 KO mouse models, we identified that NRK-2 protects the heart against DCM, cardiac fibrosis, LV chamber dilatation, and dysfunction in chronic PO conditions. Mechanistically, we identified NRK-2 as a novel regulator of the JNK MAPK pathway and our findings revealed that NRK-2 overexpression significantly attenuates JNK activation and mitochondrial dysfunction in human cardiomyocytes post-Ang II treatment. These findings highlight the protective roles of NRK-2 in PO which is partly driven by inhibiting the JNK MAPK pathway and mitochondrial function. Given the muscle-specific expression of NRK-2, inducing the underlying molecular signaling pathway through NRK-2 analog could be a potential therapeutic strategy to limit the PO-induced dilated cardiac remodeling and heart failure.

### Source of Funding

This work was supported by Competitive (1901090162-P), Targeted (1801090144-P) and COVID-19 (CoV-0302) research grants from the University of Sharjah to Firdos Ahmad. The study was also supported by an operational grant provided to the Cardiovascular Research group.

## Data Availability Statement

All the supporting data are included within the main article and its supplementary files and additional information is available from the corresponding author on reasonable request.

## Clinical perspective

∘ Despite therapeutic advancements, dilated cardiomyopathy (DCM) remains the leading cause of heart failure and death worldwide. Current therapeutic strategies are ineffective to limit DCM and remodeling and, heart transplantation remains the last line of therapy.
∘ Herein, we report the novel roles of NRK-2 in the regulation of JNK pathway and pathogenesis of dilatative cardiac remodeling. Employing preclinical *in vivo* and *in vitro* models we identified that loss of NRK-2 promotes chronic pressure overload-induced dilatative cardiac remodeling and heart failure. Consistently, NRK-2 gain-of-function attenuates angiotensin II-induced human cardiomyocyte hypertrophy, cardiomyopathy and mitochondrial dysfunction.
∘ These findings strongly suggest that gain-of NRK-2 function is protective and NRK-2 analogue could be designed and tested against pressure overload-induced dilatative cardiac remodeling and associated pathological conditions.

