## Supplementary material for "Nicotinamide riboside kinase-2 inhibits JNK pathway and limits dilated cardiomyopathy in mice with chronic pressure overload": Suppl Figures

### Suppl. Figure 1

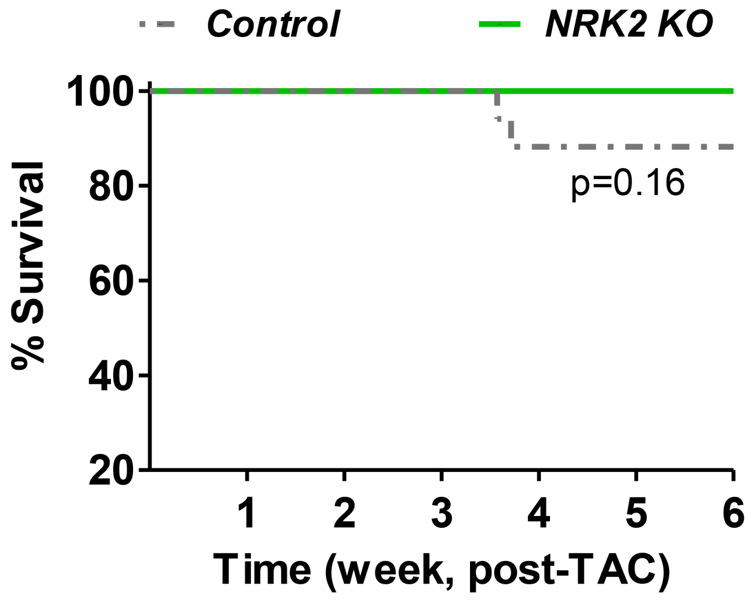


**Supplemental Figure 1.** Survival curve shows a comparable mortality between control and NRK-2 KO mice post-TAC, n=16-17. *P* value was calculated through Log-rank (Motel-Cox) test.

### Suppl. Figure 2

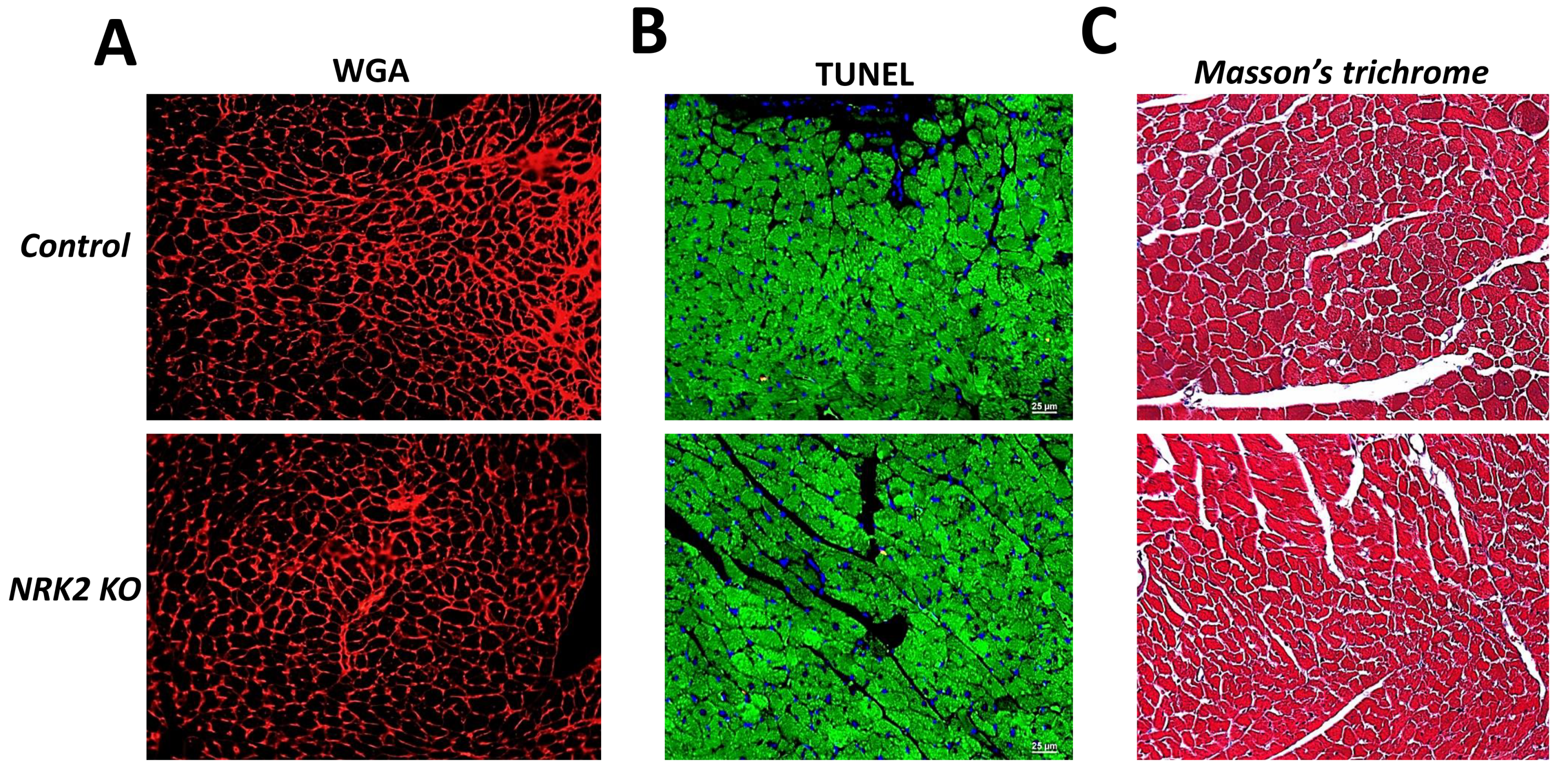


**Supplemental Figure 2.** Representative images of (**A**) wheat germ agglutinin (WGA) (**B**) TUNEL and (**C**) trichrome-stained heart sections from the sham-operated animals show a comparable cell size, apoptosis and the fibrotic area in control and NRK2 KO hearts. Images were taken using an 20X objective.
